# Mortality Risk-Stratified Septic Serum Depresses Contractility and Mitochondrial Function in Human Induced Pluripotent Stem Cell-Derived Cardiomyocytes

**DOI:** 10.64898/2025.12.21.695853

**Authors:** Andrew J. Lautz, Catherine S. Kapcar, Shuangmin Zhang, Leland Dunwoodie, Kelli Harmon, Christopher N. Mayhew, Mihir R. Atreya, Basilia Zingarelli, Genomics of Pediatric Septic Shock Investigators

## Abstract

**Background:** Sepsis-associated myocardial dysfunction (SAMD) is common in children with septic shock, is independently associated with mortality, and has no disease-modifying treatments. Differences in murine cardiomyocyte biology and repeated failures to translate discoveries into novel therapies for septic shock underscore a key translational need for human-relevant disease modeling. We sought to investigate human induced pluripotent stem cell-derived cardiomyocytes (iPSC-CMs) exposed to mortality risk-stratified septic serum as a model of SAMD.

**Methods:** Serum from children with septic shock (n=120) was stratified by Pediatric Sepsis Biomarker Risk Model (PERSEVERE) II mortality probability as low, intermediate, or high risk. We conducted aptamer-based proteomic analysis of septic serum to determine differentially expressed proteins in children with high compared to low mortality risk. We treated iPSC-CMs with risk-stratified septic serum and defined contractile and mitochondrial functional and transcriptomic responses.

**Results:** We found 612 differentially expressed proteins in children with high mortality probability, most prominently interleukins (IL)-6 and −8. High-risk septic serum reversibly depressed iPSC-CM contractility as measured by percent shortening, while low-risk septic serum had no impact. Further, high-risk septic serum depressed basal mitochondrial respiration, maximum uncoupled respiration, and coupled oxidative phosphorylation in iPSC-CMs relative to low-risk serum. We identified distinct patterns of gene expression due to risk-stratified serum with 5,293 differentially expressed genes, including upregulation of acute phase reactants and apolipoproteins and downregulation of chemokines, as well as transcriptional changes reflective of chronic IL-6 and IL-8 signaling.

**Conclusions:** Septic serum from children with high mortality risk exhibited distinct proteomic signatures, notably enriched for IL-6 and IL-8. Human iPSC-CMs differentially responded to risk-stratified septic serum, recapitulating phenotypic features of SAMD including reversible contractility depression and mitochondrial dysfunction with high-risk septic serum. These findings establish mortality risk-stratified septic serum exposure of iPSC-CMs as a human-relevant translational platform to interrogate mechanisms of myocardial dysfunction in septic shock.

## INTRODUCTION

Despite decades of research and advancements in clinical intensive care, nearly one in four children with septic shock continue to die worldwide.^1–3^ Accompanying organ dysfunctions herald poor outcomes,^4^ and sepsis-associated myocardial dysfunction (SAMD) is both common and independently associated with mortality after accounting for baseline severity of illness.^5,6^ The pathobiology of SAMD is complex,^7^ and no disease-modifying therapies have been identified, underscoring the need for more mechanistic research into cardiomyocyte dysfunction in septic shock.

Circulating myocardial depressant factors, including various cytokines, in the serum of children and adults with septic shock have been found to depress the contractile function of *in vitro* murine cardiomyocytes,^8–12^ but these models are known to differ from human cardiomyocytes in action potential, contractile protein isoforms and phosphorylation state, signaling patterns, and transcriptomes.^13,14^ To date, few molecular discoveries from animal studies have translated into successful clinical trials in human septic shock,^15–18^ necessitating human modeling of cardiomyocyte dysfunction. Human induced pluripotent stem cells (iPSCs) are a renewable source of human cardiomyocytes through small molecule-based differentiation^19,20^ and have been used to model viral myocarditis and doxorubicin cardiotoxicity and for high-throughput drug screening.^21–24^ We sought to investigate the human iPSC-cardiomyocyte (iPSC-CM) response to human septic serum as a translational model of SAMD.

We previously analyzed the whole blood transcriptome from children with septic shock, identifying differences in gene expression in children with high versus low mortality probability as classified by the Pediatric Sepsis Biomarker Risk Model (PERSEVERE)-II,^25,26^ linking dysregulated neutrophil-driven inflammation with poorer outcomes.^27^ Here, we first investigated the serum proteome of children with septic shock, stratifying risk by PERSEVERE II, given the association of PERSEVERE biomarkers with pediatric SAMD^28^ and to better understand serum factors associated with mortality as a prelude to experimental serum exposure to iPSC-CMs. We delineated a distinct proteomic shift moving from low to intermediate to high PERSEVERE mortality risk and distinguished interleukins (IL)-6 and −8 as the most differentially expressed proteins between high- and low-risk serum. Next, we subjected iPSC-CMs to risk-stratified serum, detailing changes in contractility, respirometry, and gene expression. We employed metabolic maturation through induced fatty acid oxidation^29^ to mitigate issues with the immature phenotype of iPSC-CMs^30^ and subjected iPSC-CM spheroids to mortality risk-stratified serum banked from children with septic shock. We demonstrated that exposure to high-risk serum depressed iPSC-CM contractility, as well as basal and maximum uncoupled respiration and coupled oxidative phosphorylation relative to low-risk septic serum. Bulk RNA sequencing revealed distinct patterns of gene expression in iPSC-CMs responding to high versus low-risk septic serum, including downstream alterations related to IL-6 and IL-8 signaling.

## METHODS

These studies employed serum samples from the Sepsis Genomics Collaborative, an ongoing, prospective, multi-center observational cohort of pediatric septic shock, which has been detailed previously.^31–33^ All study procedures involving human subjects were approved by the Institutional Review Board (IRB) at Cincinnati Children’s Hospital Medical Center (CCHMC), serving as the central IRB, and by the local IRBs at participating institutions; these studies were conducted in accordance with the Declaration of Helsinki and its subsequent amendments or comparable ethical standards. Blood was previously collected and biobanked after written informed consent from parents or legal guardians (IRBs 2008-0558 and 2022-0721), and donor iPSCs were likewise previously developed and banked by the CCHMC Pluripotent Stem Cell Facility (PSCF) after written informed consent (IRB 2017-2011) from parents or legal guardians. The de-identified RNA sequencing and proteomic data in this study have been deposited in the NCBI Gene Expression Omnibus and will be publicly accessible upon publication.

### Serum Collection and Mortality Risk Stratification

Blood collection and mortality risk stratification using Pediatric Sepsis Biomarker Risk Model (PERSEVERE) II occurred as previously described.^26,28^ Briefly, children less than 18 years of age who met consensus criteria for septic shock^34^ were enrolled after informed consent was obtained from their parents or legal guardians from 13 pediatric intensive care units across the United States from 2006 to 2021. Blood samples were obtained within 24 hours of enrollment, and clinical laboratory data were collected for PERSEVERE II determination for estimation of baseline mortality probability.^25,26^ The PERSEVERE biomarkers include IL-8, C-C chemokine ligand 3, matrix metallopeptidase 8, heat shock protein 70kDa 1B (HSPA1B), and granzyme B, and concentrations were measured using a multiplex magnetic bead platform (MILLIPLEX MAP) designed for this project by the EMD Millipore Corporation (MilliporeSigma) and a Lumine× 100/200 System (Luminex Corporation) as previously published.^35^ Serum biomarker concentrations and the admission platelet count were used to calculate PERSEVERE II baseline mortality probability, ranging from 0.000 to 0.571.^25^ Patients were stratified as being low risk (≤1.9%), intermediate risk (16.7-18.9%), and high risk (≥30%) of mortality as previously described^28^ for proteomic analysis. Downstream serum exposure of iPSC-CMs only utilized high- and low-risk serum samples to maximize pathobiological differences across the spectrum of pediatric septic shock (**Figure 1a**).

**Figure 1:**
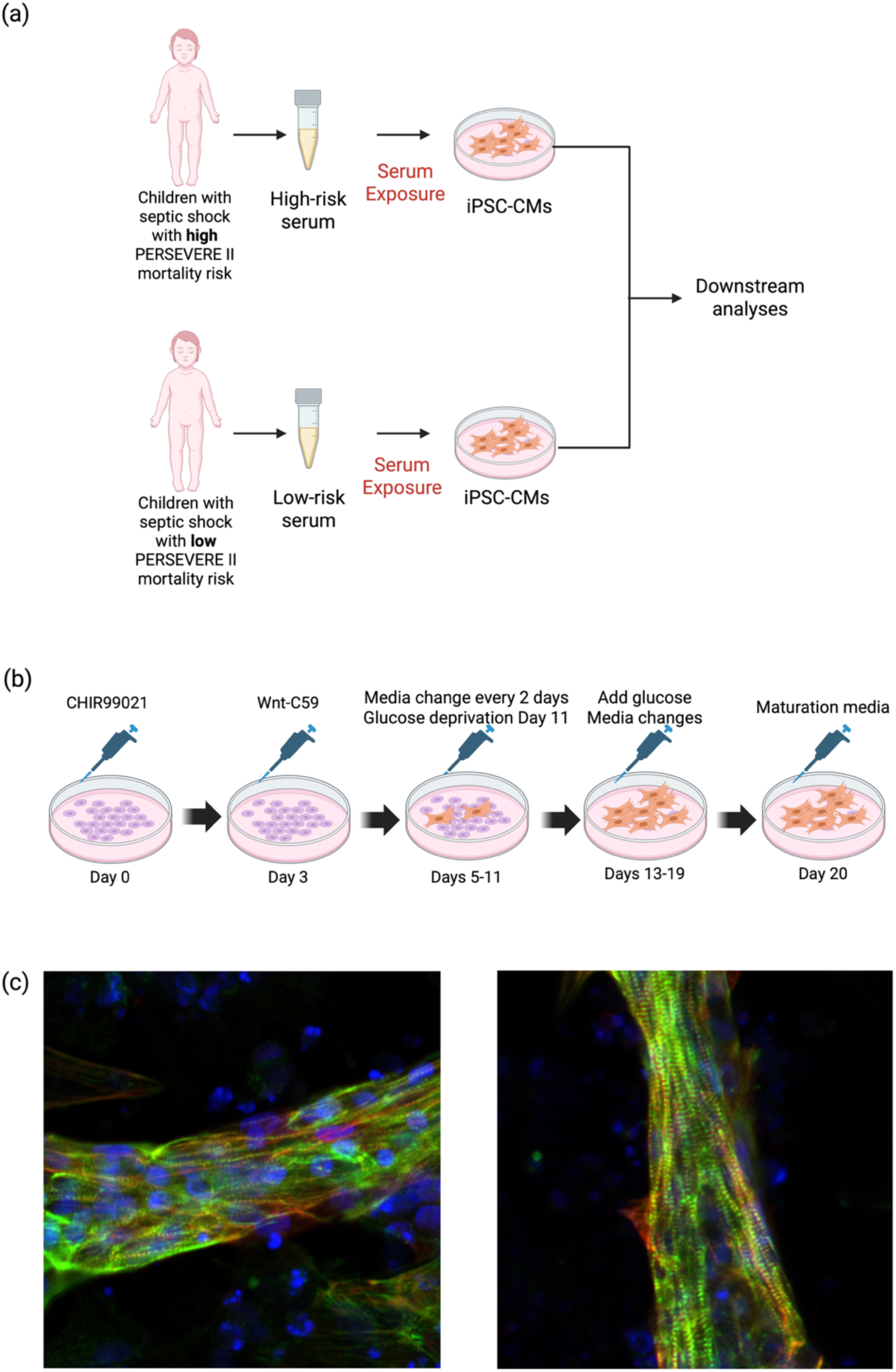
Differentiation of iPSC-CM spheroids. (**a**) Schematic representation of septic serum risk stratification and exposure to iPSC-CMs, and (**b**) schematic representation of iPSC-CM differentiation. Created with BioRender.com. (**c**) Characterization of iPSC-CMs by immunofluorescence. iPSC-CMs imaged at 20 days (left panel) and 39 days (right panel) after differentiation. Blue = DAPI, Green = alpha-actinin, Red = cardiac troponin T.

### Proteomic Analyses

Serum samples banked from children with septic shock were analyzed with the SomaScan 7k Assay (SomaLogic, Inc.). Raw ADAT files were read with the R package SomaDataIO, and samples were integrated with metadata including PERSEVERE risk category (n=76 low risk, n=23 intermediate risk, and n=21 high risk), age, and study site. Plate identifiers were read and treated as a batch variable for adjustment in downstream analyses. After removing expression data for internal controls, there were 7,280 human protein epitopes with measurements of relative fluorescence units, which were log_2_ transformed for analysis. Expression matrices adjusted for plate, age, and site were created using limma and the removeBatchEffect function. Principal component analysis (PCA) was performed on adjusted matrices using the top 1,000 most variable proteins, generating two-dimensional (principal component 1, PC1, to PC2; PC2 to PC3) plots and a three-dimensional PCA plot with ellipsoids drawn using 68% confidence intervals for each PERSEVERE II risk category. Intermediate-risk samples were included in PCA and subsequent heatmap visualizations but excluded from inferential analyses with the comparison of interest being high versus low risk.

Differential protein expression analysis was conducted using linear models with empirical Bayes moderation in the R package limma, comparing high-risk to low-risk patients. Expression matrices adjusted for plate, age, and site were analyzed with lmFit with empiric Bayes variance moderation with robust estimation, and differentially expressed proteins (DEPs) were summarized with log_2_ fold change and identified as statistically significant at a Benjamini-Hochberg adjusted p-value < 0.05. Permutational multivariate analysis of variance (PERMANOVA) assessed global differences in the high- and low-risk proteome using the top 1,000 most variable proteins and Euclidean distance matrices with 999 permutations. Beta dispersion testing was performed using the betadisper function in the R package vegan with permutation testing to determine whether PERMANOVA results also reflected differences in within-group dispersion.

Differential expression results were visualized with a volcano plot with significance thresholds set at log_2_ fold change ≥0.58 (∼1.5-fold change) and false discovery rate (FDR) <0.05 and with a heatmap constructed using the top 50 differentially expressed proteins adjusted for plate, age, and site and z-score scaled across samples. Functional enrichment analyses of significantly up- and down-regulated proteins using Reactome and Gene Ontology Biological Process pathways were performed, and Benjamini-Hochberg adjusted pathway p-values < 0.05 were considered significant.

### Differentiation of iPSC-CMs

Healthy donor iPSCs were obtained as a clonal line from the CCHMC PSCF biobank plated onto Matrigel (Corning, cat. no. 354234) or Cultrex SCQ (Biotechne, cat. no. 3434-010-02)-coated 6-well and 24-well plates (at densities of 1×10^6^ cells per well for 6-well plates and 2×10^5^ cells per well for 24-well plates). We adapted an established method for chemically differentiating iPSC-CMs from human iPSCs using small molecules and metabolic purification,^19,36^ and modified the protocol to culture in maturation media to improve electrophysiologic and mechanical maturity^29^ (**Figure 1b**). Cells were maintained in mTeSR1 (StemCell Technologies, cat. no. 85851) medium changed daily until differentiation started when the cells were at approximately 90-95% confluence by visual examination.

Upon reaching near confluence (day 0), iPSC media was changed to RPMI 1640 (Thermo Fisher Scientific, cat. no. 11875093) with 1x B27 minus insulin (Thermo Fisher Scientific, cat. no. A1895601) treated with 4uM CHIR99021 (Tocris, cat. no. 4423) for 3 days. Subsequently (day 3) the cells were treated with the 2uM Wnt inhibitor C59 (Tocris, cat. no. 5148) in RPMI1640 with B27 minus insulin for another 2 days. Media was changed with fresh RPMI1640 with B27 minus insulin every 2 days until day 11 when differentiating cells were changed to RPMI1640 without glucose (Thermo Fisher Scientific, cat. no. 11875020) with B27 plus insulin (Thermo Fisher Scientific, cat. no. 12587010) and 5mM sodium L-lactate (Sigma Aldrich, cat. no. 71718) for metabolic enrichment through transient glucose deprivation.^19,37^ Glucose was added back after 2 days with RPMI1640 (Thermo Fisher Scientific, cat. no. 11875093) with B27 plus insulin, and media was changed every 2-3 days until day 20, when cells were transitioned to maturation media rich in oxidative substrates and low in glucose.^29^

Maturation media was composed of DMEM without glucose (Thermo Fisher Scientific, cat. no. 11966025) supplemented with 5mM sodium L-lactate (Sigma Aldrich, cat. no. 71718), 3mM D-glucose (Sigma Aldrich, cat. no. G7021), 10mM L-lactate (Sigma Aldrich, cat. no. 71718), 5mg/ml vitamin B12 (Sigma Aldrich, cat. no. V6629), 0.82mM biotin (Sigma Aldrich, cat. no. B4639), 5mM creatine monohydrate (Sigma Aldrich, cat. no. C3630), 2mM taurine (Sigma Aldrich, cat. no. T0625), 2mM L-carnitine (Sigma Aldrich, cat. no. C0283), 0.5mM ascorbic acid (Sigma Aldrich, cat. no. A8960), 1x NEAA (Thermo Fisher Scientific, cat. no. 11140), 0.5% (w/v) Albumax (Thermo Fisher Scientific, cat. no. 11020021), 1x B27 plus insulin (Thermo Fisher Scientific, cat. no. 12587010), and 1% KOSR (Thermo Fisher Scientific, cat. no. 10828028). Spontaneously beating iPSC-CM spheroids were typically evident by day 14 of differentiation and continued through at least 75 days post-differentiation. Analyses were conducted on iPSC-CM spheroids at least 35 days post-differentiation, reflecting more than two weeks exposure to maturation media.

### Immunofluorescence Analysis

Cells were rinsed with phosphate buffered saline with Tween (PBST) and fixed in 4% paraformaldehyde in phosphate buffered saline (PBS) for 15 minutes at room temperature. After 3 washes in PBST (5 minutes each), cells were permeabilized in 0.1% saponin (Sigma Aldrich, cat. no. S7900) in PBS for 20 minutes at room temperature followed by incubation in blocking solution with 4% goat serum (Sigma Aldrich, cat. no. G9023) and 4% donkey serum (Sigma Aldrich, cat. no. D9663) in PBS for 60 minutes at room temperature. The following primary antibodies were diluted in 0.1% saponin in PBS: alpha-actinin produced in mouse at 1:800 (Sigma Aldrich, cat. no. A7811) and cardiac troponin T produced in rabbit at 1:400 (Abcam, cat. no. ab45932). Samples were incubated on a shaker overnight at 4 degrees Celsius, rinsed 3 times in PBST (5 minutes each) and incubated in secondary antibodies (Alexa 488 conjugated to goat anti-mouse IgG1, Invitrogen cat. no. A21121, and Alexa 568 conjugated to donkey anti-rabbit IgG polyclonal, Invitrogen cat. no. A10042) diluted in 0.1% saponin in PBS at 1:200 in the dark for 2-3 hours. Cells were then washed 3 times in PBST (5 minutes each) and incubated with DAPI (Invitrogen, cat. no. D1306) at 1:1000 diluted in PBS for 5 minutes. Cells were rinsed a final time in PBST for 5 minutes and kept in PBS for image capture. Images were obtained with the Nikon Eclipse Ti2 microscope.

### Contractility Measurement

An IonOptix Cardiomyocyte Calcium and Contractility System with an Inverted Olympus IX73 Fluorescence Microscope and CytoMotion software (IonOptix) was utilized to measure pixel-based, real-time contractility of iPSC-CM spheroids in 6-well and 24-well cell culture plates. Contractility traces were measured using IonWizard (version 7.8) and analyzed with CytoSolver (version 3.0.2). Primary measurements of contractility were percent shortening, as determined by pixel correlation, and spontaneous beat frequency. Repeated measures were conducted after documenting spheroid location within each well of the culture plate and capturing baseline images to align the region of interest in IonWizard. To verify reliability of repeated measurements, we measured iPSC-CM spheroids in standard maturation media serially on three separate days with 24 hours between each measurement. There was no difference between measurements of percent shortening within individual spheroids (p=0.18) by repeated measures analysis of variance (**Supplemental Figure 1**), validating this approach to contractility assessment.

### Mitochondrial Respirometry

The Seahorse XFe24 Analyzer (Agilent) was used to assess mitochondrial respirometry using the Seahorse XF Mito Stress Test Kit (Agilent). Seahorse XF24 cell culture microplates (Agilent) were coated with Matrigel or Cultrex SCQ the day prior to plating iPSC-CMs. On the day of replating, iPSC-CMs were washed with pre-warmed PBS (37 degrees Celsius; 1mL per well for 24-well plates, 3mL per well for 6-well plates) and treated with pre-warmed STEMdiff Cardiomyocyte Dissociation Medium (StemCell Technologies, cat. no. 05026; 500μL per well for 24-well plates, 1mL per well for 6-well plates) at 37 degrees Celsius for 7-10 minutes. Pre-warmed STEMdiff Cardiomyocyte Support Medium (StemCell Technologies, cat. no. 05027; 500μL per well for 24-well plates, 1mL per well for 6-well plates) was added and pipetted up and down gently to detach the cells. Cells were collected in a conical tube and counted (20μL mixed with 20 μL Trypan blue) and then centrifuged at 200g for 5 minutes at room temperature. The supernatant was discarded, and cells were resuspended in STEMdiff Cardiomyocyte Support Medium at a concentration of 1.5×10^6^ cells/mL and seeded at a density of 1.5×10^5^ cells per well (100μL). Cells were incubated at 37 degrees Celsius, and an additional 150μL of medium was added one hour later before incubating overnight. The subsequent day the cells were changed to 250μL maturation media in each well and incubated an additional 24-48 hours prior to serum exposure. After 24 hours exposure to 10% risk-stratified septic serum in media, cells underwent respirometry.

One hour prior to the assay, the old medium was removed, and 500uL assay medium (Seahorse XF DMEM Medium, Agilent cat. no. 103575-100) supplemented with 10mM glucose (Sigma Aldrich, cat. no. G7021, 2mM glutamate (Thermo Fisher Scientific, cat. no. J60573.14), and 1mM pyruvate (Sigma Aldrich, cat. no. P2256) was added to each well. The cells were then incubated at 37 degrees Celsius in a non-CO_2_ incubator for one hour until loading into the Seahorse XFe24 Analyzer. As part of the Mito Stress Test Kit, sequential injections were oligomycin at 1.5 μM, FCCP at 0.5 μM, and rotenone/antimycin A at 0.5 μM. The oxygen consumption rates were calculated using the Seahorse Wave controller Software 2.6.1 (Agilent). After the experiment cell counts were calculated from each well (20μL samples mixed with 20μL Trypan Blue), and data were normalized to live cell counts per well using Microsoft Excel for Mac (version 16.103, Microsoft).

### Bulk RNA Sequencing

After iPSC-CMs were exposed for 24 hours to 10% mortality risk-stratified septic serum diluted in media, RNA was isolated with the miRNeasy Micro Kit (Qiagen, cat. no. 217084). Bulk RNA was sequenced in conjunction with the CCHMC Genomics Sequencing Facility. RNA (150 to 300 ng as determined by Invitrogen^TM^ Qubit^TM^ high-sensitivity spectrofluorometric measurement) was poly-A selected and reverse transcribed using the Illumina Stranded mRNA Prep, Ligation (cat. no. 20040534) and Index Plate Set A. Each sample was fitted with one of 96 adapters containing a different 8 base molecular barcode for high-level multiplexing. After 15 cycles of polymerase chain reaction amplification, completed libraries were sequenced on an Illumina NovaSeq^TM^ 6000, generating 50 million high-quality 150 base-long paired-end reads per sample. Fastq files were subjected to quality control to assess the need for trimming of adapter sequences or bad quality segments, using FastQC v0.11.7, Trim Galore! v0.4.2, and cutadapt v1.9.1. The trimmed reads were aligned to the reference human genome version hg38 with the program STAR v2.6.1e. Aligned reads were stripped of duplicate reads with the program sambamba v0.6.8. Gene-level expression was assessed by counting features for each gene, as defined in the NCBI’s RefSeq database. Read counting was done with the program featureCounts v1.6.2 from the Rsubread package. Raw counts were normalized as transcripts per million. Differentially expressed genes (DEGs) between groups were assessed with the R package DESeq2 v1.26.0. Gene list and log2 fold change were used for Gene Set Enrichment Analysis (GSEA) using Gene Ontology (GO) pathway datasets. Plots were generated using the ggplot2 package and base graphics in R.

### Statistical Analyses

For proteomic analysis, DEPs were identified at a Benjamini-Hochberg adjusted p-value < 0.05. Contractility data (percent shortening and spontaneous beat frequency) are presented in text as median with interquartile range and analyzed with Wilcoxon matched-pairs signed rank tests for comparison to baseline. Repeated measures analysis of variance (ANOVA) was employed for repeated measures analysis of individual iPSC-CM spheroids as delineated in text. Respirometry measurements of oxygen consumption rate normalized to the number of live cells are presented in text as mean ± standard error of the mean and compared with unpaired t tests. For RNA sequencing analysis, DEGs were defined as a log_2_ fold change ≥ |0.58| (∼1.5-fold change) and a false discovery rate (FDR) q-value < 0.05. Normalized enrichment scores (NES) for GSEA with an FDR q-value < 0.05 were considered statistically significant. The number (n) of data points per experiment is delineated in the figure legends. Statistical tests were performed with GraphPad Prism 10 Software.

## RESULTS

### Serum proteomics revealed IL-6 and IL-8 as key differentially expressed proteins in high mortality-risk septic serum

We first sought to define the proteomic differences in mortality risk-stratified septic serum (n=120) as classified by PERSEVERE II as a foundation for modeling SAMD with serum exposure to iPSC-CMs. We observed a clear shift in the serum proteome moving from low to intermediate to high mortality risk (**Figure 2a**). After adjustment for batch effects, age, and site, PERSEVERE II mortality risk (high versus low risk) remained significantly associated with proteomic composition, accounting for 5.3% of the variance across all proteins (PERMANOVA R^2^ = 0.053, p=0.008) and 7.0% of the variance among the top 1000 most variable proteins (R^2^ = 0.070, p=0.003). This was accompanied by an increase in proteomic heterogeneity in high-risk serum (permutation testing p=0.002 for all proteins and p=0.005 for the top 1000 most variable proteins), suggesting both a shift in centroid separation and differences in dispersion (**Supplemental Table 1**), reflecting the clinical heterogeneity of human septic shock. We identified 612 differentially expressed proteins comparing children with high mortality probability to children with low mortality probability after adjusting for batch effects, age, and site. Heatmapping revealed discrete patterns in the top 50 DEPs among low to intermediate to high mortality risk (**Figure 2b**). IL-6, identified by two aptamers (log_2_ fold change 3.16, adj p=1.2×10^−6^; and log_2_ fold change 2.94, adj p=2.9×10^−6^) and IL-8 (CXCL8, log_2_ fold change 2.97, adj p=6.5×10^−9^) were among the most differentially expressed proteins among pediatric septic shock patients with high mortality probability (**Figure 2c** and **Supplemental Table 1**). Enrichment analyses using Reactome and GO Biological Process annotations delineated distinct pathways over- and under-represented in high-risk patients, including over-representation of the Reactome pathway Signaling by Interleukins in children with high mortality probability (**Figure 2d** and **Supplemental Figure 2**). These data define the differences in high-and low-risk serum exposure to iPSC-CMs and highlight the potential role for IL-6 and IL-8 signaling in the cardiomyocyte response to septic serum.

**Figure 2:**
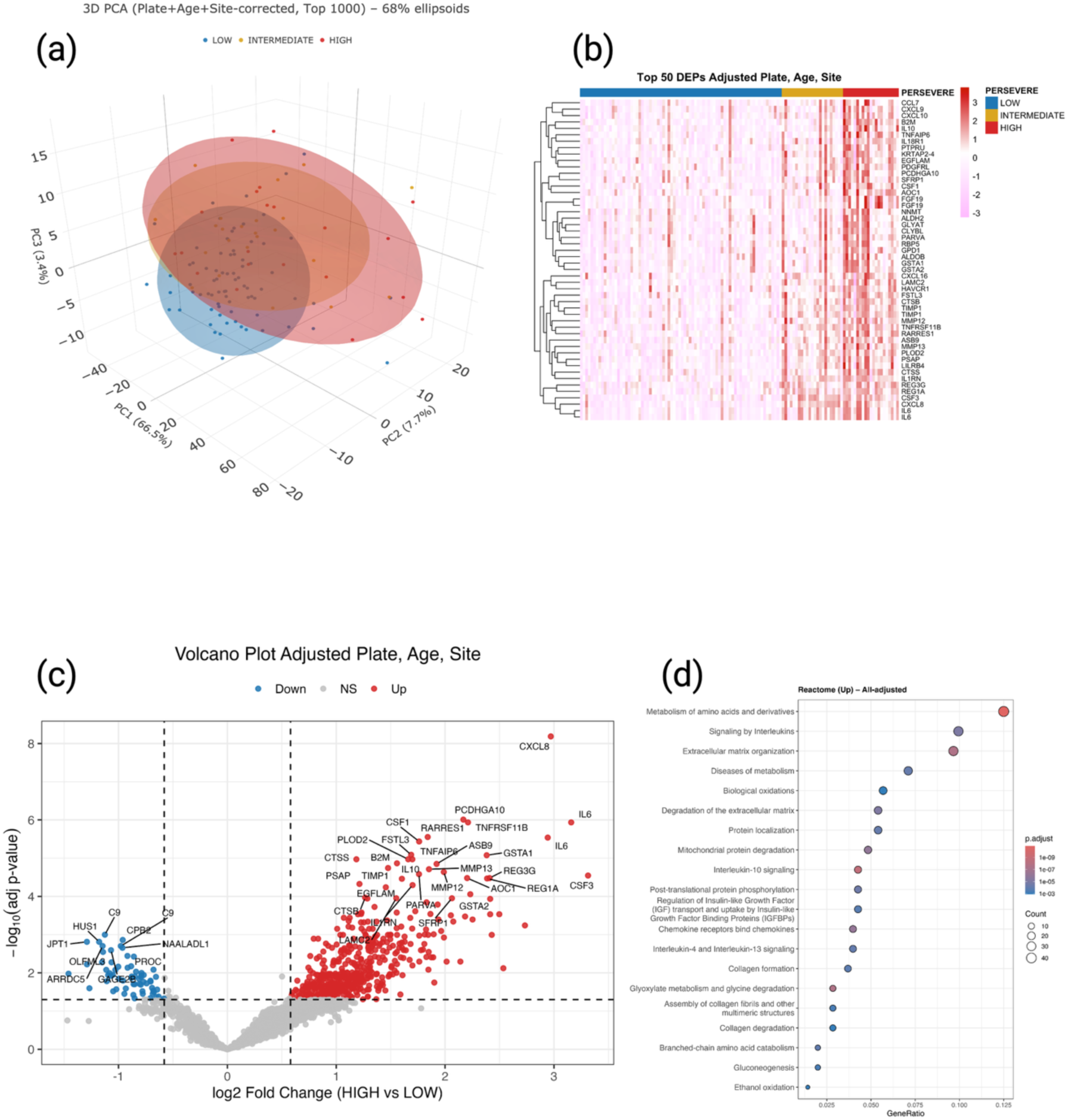
High mortality risk shifts the serum proteome and is associated with high serum levels of IL-6 and IL-8. Three-dimensional Principal Component Analysis plot of serum from children with low PERSEVERE II mortality risk (blue, n=76), intermediate PERSEVERE II mortality risk (yellow, n=23), and high PERSEVERE II mortality risk (red, n=21). (**b**) Heatmap of top 50 differentially expressed proteins between high- and low-risk samples, adjusted for plate, age, and site, visualized across low-, intermediate-, and high-risk strata. (**c**) Volcano plot of differentially expressed proteins in high-risk children with septic shock compared to low-risk children with septic shock using log_2_ fold change >= |0.58| and adjusted p-value < 0.05, after adjusting for plate, age, and site. IL-6 is identified by two aptamers. CXCL8 = IL-8. (**d**) Reactome pathway annotation of over-represented proteins among children with high mortality probability with a Benjamini-Hochberg adjusted p-value < 0.05 after adjustment for plate, age, and site.

### High mortality-risk septic serum reversibly depressed iPSC-CM contractility

We differentiated iPSC-CMs that durably expressed alpha-actinin and cardiac troponin T in an organized substructure as imaged by immunofluorescence at two different time points during differentiation (**Figure 1c**). iPSC-CM spheroids exhibited regular spontaneous contractions (**Supplemental Figure 3**) that were quantified with pixel-based real-time tracking using the IonOptix system. Exposure to high mortality-risk serum banked from children with septic shock (10% serum in medium) depressed both spontaneous beat frequency [0.20 Hz (0.15-0.27 Hz) vs. baseline 0.33 Hz (0.22-0.41 Hz), p=0.01] and percent shortening [14.4% (2.5-30.1%) vs. baseline 23.2% (16.7-30.6%), p=0.03] at 24 hours. In contrast, iPSC-CMs subjected to low-risk serum demonstrated no changes in spontaneous beat frequency [0.30 Hz (0.27-0.39 Hz) vs. baseline 0.26 Hz (0.22-0.34 Hz), p=0.52] or percent shortening [22.1% (10.8-33.3%) vs. baseline 25.7% (17.0-38.3%), p=0.43] (**Figure 3a**). In separate experiments focused on reversibility, iPSC-CMs exposed to high mortality-risk serum exhibited lack of any spontaneous beating at 24 hours, while low-risk serum did not affect beat frequency [0.21 Hz (0.17-0.26 Hz) vs. baseline 0.23 Hz (0.18-0.25 Hz), p>0.99] nor contractility [22.5% (17.9-25.9%) vs. baseline 19.6% (9.7-29.0%), p=0.48]. After 20 hours of recovery in medium without septic serum, the high-risk group resumed spontaneous contractility (**Figure 3b**), confirming reversibility of the phenotype consistent with clinical SAMD. Using repeated measures of individual iPSC-CM spheroids prior to and one hour after exposure to risk-stratified septic serum, there was no significant difference in percent shortening after low-risk exposure (p=0.80), while high-risk serum significantly depressed percent shortening (p=0.01), further validating the aggregate data demonstrating differential response to mortality risk-stratified septic serum (**Figure 3c**).

**Figure 3:**
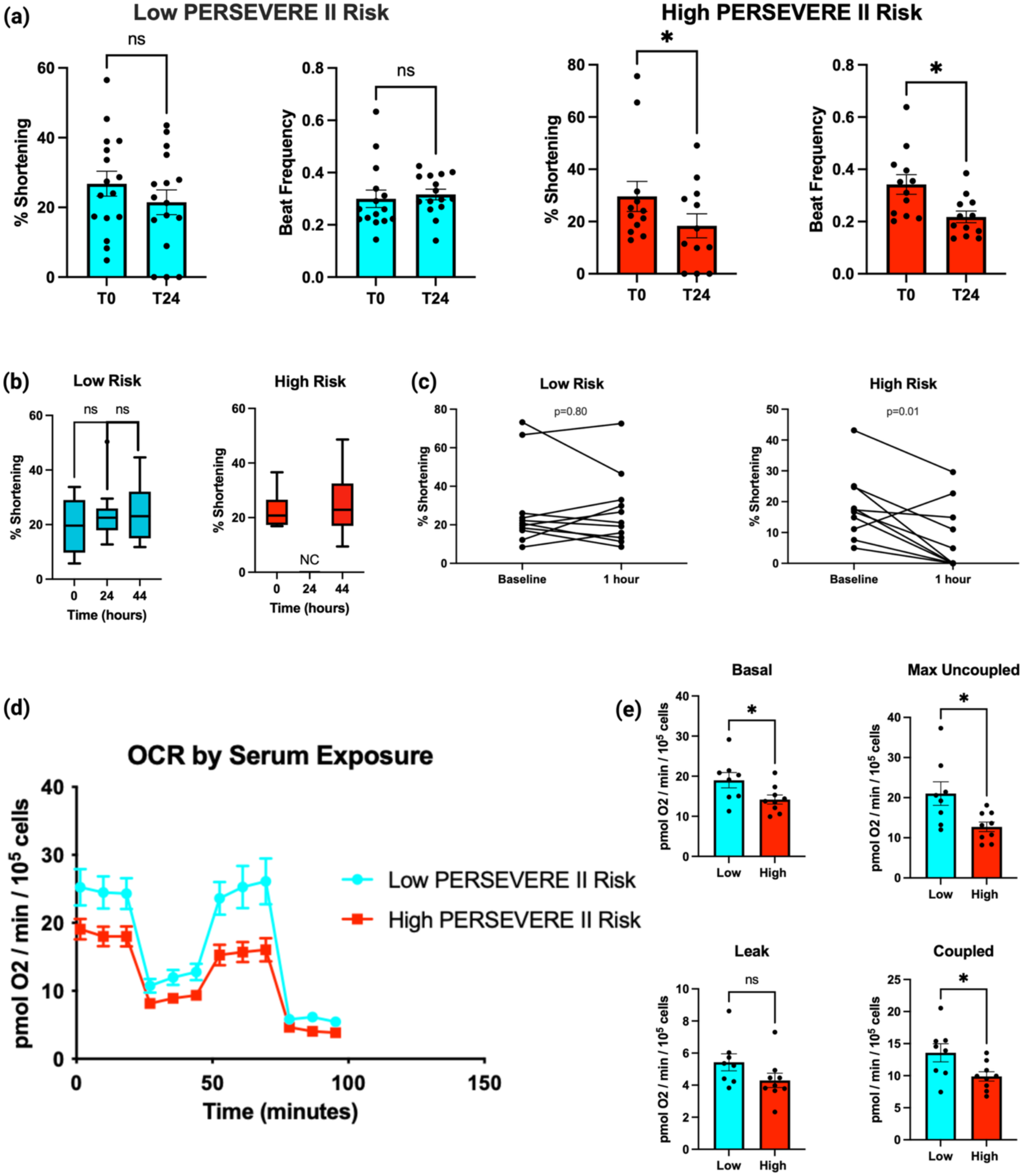
High mortality-risk septic serum reversibly depressed iPSC-CM contractility and depressed mitochondrial respiration relative to low-risk serum. (**a**) Percent shortening and beat frequency of iPSC-CMs before and 24 hours after 10% low-risk (left panel) and high-risk (right panel) septic serum exposure. n=12-16 per group. (**b**) Percent shortening of iPSC-CMs exposed to 7.5% septic serum from children at either low-risk (left panel) or high-risk (right panel) of mortality. NC = not contracting. n=9-12 group. (**c**) Repeated measurements of percent shortening of individual iPSC-CM spheroids before and one hour after exposure to low-risk (left panel) and high-risk (right panel) septic serum. n=10-11 per group. (**d**) High resolution respirometry of iPSC-CMs exposed to mortality risk-stratified septic serum with oxygen consumption rate (OCR) over time with sequential injection of mitochondrial inhibitors and uncouplers. (**e**) Bioenergetics quantified as differences in basal respiration, maximum uncoupled respiration, leak respiration, and coupled oxidative phosphorylation after exposure to mortality risk-stratified septic serum. n=8-9 per group.

### High mortality-risk septic serum depressed iPSC-CM mitochondrial respiration

Risk-stratified septic serum also differentially impacted iPSC-CM mitochondrial respiration. After 24 hours exposure to 10% high-risk septic serum in medium, iPSC-CM spheroids exhibited diminished basal respiration [14.2 ± 1.1 vs. 19.0 ± 1.9 pmol O_2_/min/10^5^ cells, p=0.04], maximum uncoupled respiration (12.7 ± 1.2 vs. 21.0 ± 2.9, p=0.02), and coupled oxidative phosphorylation (9.9 ± 0.7 vs. 13.6 ± 1.4, p=0.03) without changing leak respiration (4.3 ± 0.4 vs. 5.4 ± 0.5, p=0.12) compared to low-risk septic serum (**Figure 3d**). These findings, normalized to the number of live cells per well, suggest widespread changes in mitochondrial respiration induced specifically by septic serum from high-risk patients relative to serum from patients with sepsis but low mortality probability.

### Gene expression in iPSC-CMs was altered with high-risk septic serum

Similarly, 24 hours of exposure to 10% risk-stratified septic serum led to significant differences in the iPSC-CM spheroid transcriptome. There were 5,293 DEGs in iPSC-CMs treated with high-risk serum relative to low-risk septic serum (**Supplemental Table 2**). Principal Component Analysis revealed separate clustering of high- and low-risk groups (**Figure 4a**), and we identified distinct patterns of gene upregulation and downregulation between risk strata (**Figure 4b** and **4c**). GSEA delineated 154 GO pathways upregulated and 11 GO pathways downregulated with high-risk serum with q-value <0.05 (**Supplemental Table 3**). Upregulated GO pathways (**Figure 4d**) with high-risk serum treatment included Acute Phase Response (NES 2.5, q-value 0.0), with four of the overall top ten up-regulated DEGs: haptoglobin (HP), alpha 2-HS glycoprotein (AHSG), orosomucoid (ORM1), and lipopolysaccharide binding protein (LBP), as well as interleukin-6 receptor (IL6R); and Abnormality of Lipoprotein Cholesterol Concentration (NES 2.4, q-value 0.0), with two of the overall top ten up-regulated DEGs: apolipoprotein C3 (APOC3) and albumin (ALB), as well as multiple other apolipoproteins (APOA1, APOA2, APOA5, APOB, and APOC2). Downregulated GO pathways (**Figure 4e**) in the high-risk group included Chemokine Receptor Binding (NES −2.6, q-value 0.0) with multiple chemokine transcripts, including IL-8.

**Figure 4:**
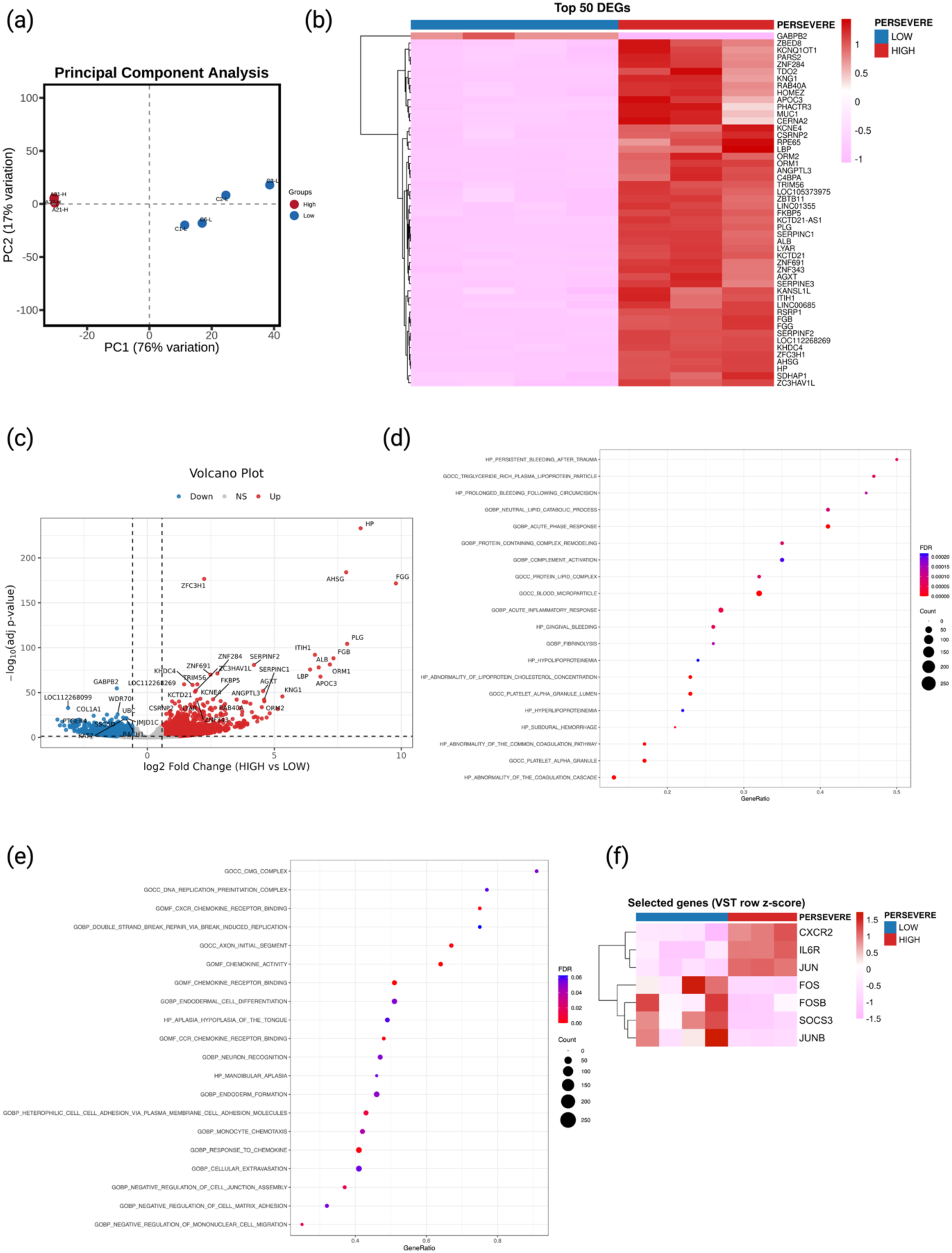
Risk-stratified septic serum distinctly alters iPSC-CM gene expression. (**a**) Principal Component Analysis plot of iPSC-CMs exposed to high-risk (n=3) and low-risk (n=4) septic serum. (**b**) Heatmap of top 50 differentially expressed genes and (**c**) volcano plot of iPSC-CMs subjected to high-versus low-risk serum using log_2_ fold change >= |0.58| and q-value < 0.05. Gene Ontology pathways upregulated (**d**) and downregulated (**e**) in iPSC-CMs exposed to high-risk serum compared to low-risk septic serum. (**f**) Heatmap of variance-stabilizing transformation (VST) expression of select genes of biologic interest, scaled per gene (row z-scored) to highlight relative differences across samples.

Further, the iPSC-CM RNA sequencing data obtained after 24 hours of high-risk serum exhibited features consistent with chronic exposure to IL-6 and IL-8 (**Figure 4f**). In addition to the acute phase response, high-risk serum was associated with upregulation of the IL-6 receptor (IL6R, log_2_ fold change 1.95, adj p=4.2×10^−10^) and downregulation of SOCS3 (log_2_ fold change −1.65, adj p=6×10^−9^) as well as upregulation of IL-8 receptor CXCR2 (log_2_ fold change 2.43, adj p=3.1×10^−7^). Downstream changes were evident in downregulation of FOS (log_2_ fold change −1.25, adj p=1.4×10^−3^), FOSB (log_2_ fold change −1.21, adj p=9.2×10^−3^), JUNB (log_2_ fold change −0.66, adj p=2.7×10^−2^), although JUN (log_2_ fold change 0.75, adj p=8.9×10^−13^) was upregulated.

## DISCUSSION

Sepsis-associated myocardial dysfunction has no targeted therapeutic treatments. Our data suggest that human cardiomyocytes differentiated from induced pluripotent stem cells can be leveraged to study the pathobiology underlying the response to the sepsis microenvironment. We established proteomic shifts in serum from children with septic shock among low-, intermediate-, and high-risk strata as estimated by PERSEVERE II, and we demonstrated that iPSC-CMs exhibit distinctive biologic responses to risk-stratified septic serum. While exposure to low-risk serum banked from children with septic shock did not alter iPSC-CM contractility, high-risk serum depressed measures of contractility in a reversible manner consistent with the clinical course of sepsis-associated myocardial dysfunction.^7,38,39^ Similarly and as seen in SAMD, high-risk serum depressed mitochondrial respiration and changed the transcriptome in iPSC-CMs relative to septic serum stratified as low risk.^40–43^ While prior models have subjected iPSC-CMs to endotoxin,^44^ this novel application of risk-stratified septic serum facilitates investigation of the cardiomyocyte response to a severe sepsis phenotype associated with high risk of mortality. Human cardiomyocyte modeling of SAMD closes an important translational gap as a necessary step toward identifying and testing potential therapeutic targets. While iPSC-CMs are known to exhibit immaturity in electrophysiology, contractility, and metabolism,^30,45^ our approach employs metabolic maturation with fatty acid oxidation to improve the fidelity of SAMD modeling as has been seen in other cardiomyocyte disease processes.^29^

We identified IL-6 and IL-8 as among the most differentially expressed proteins in the serum of children with high PERSEVERE II mortality probability compared to children with sepsis but low PERSEVERE II mortality probability. Interleukin-6 has been linked to myocardial dysfunction in meningococcal septic shock in children^46^ and has been known to depress contractility *in vitro* using animal cardiomyocyte models.^47,48^ It is expected to see IL-8 differentially expressed in high-risk patients, since IL-8 is one of the PERSEVERE II biomarkers. However, IL-8 levels have been associated with myocardial dysfunction in septic adults,^39^ and human serum with high IL-8 concentration has been demonstrated to depress cardiomyocyte contractility *in vitro*.^49,50^ The pathobiologic roles of IL-6 and IL-8 in SAMD merit further research.

Although not directly assessed in the present study, the iPSC-CM response to high-risk serum is consistent with effects of IL-6 and IL-8 signaling. RNA sequencing data identified an iPSC-CM acute phase response, a process classically driven by IL-6 signaling,^51,52^ as well as a potentially compensatory downregulation and feedback inhibition in chemokine transcription after 24 hours exposure to high-risk serum. In addition to effects on contractility, IL-6 has been identified to suppress mitochondrial respiration via JAK-STAT1/3 signaling with decreases in basal respiration, maximal uncoupled respiration, and coupled oxidative phosphorylation in a manner similar our study.^53^ It is interesting to note that these changes in iPSC-CM mitochondrial respiration with high-risk serum were also associated with an increase in multiple apolipoprotein transcripts. Whether this reflects modulation of lipid bioavailability and fatty acid oxidation or mediation of mitochondrial injury, as with APOC3 in macrophages,^54^ requires further investigation.

After 24 hours of high-risk septic serum, iPSC-CMs exhibited upregulation of IL6R and CXCR2 yet downregulation of SOCS3, a key negative feedback inhibitor of gp130/JAK/STAT signaling.^55,56^ Although SOCS3 expression is normally rapidly induced by IL-6,^57^ some temporal evidence suggests a return to baseline within 90 minutes,^58^ and chronic IL-6 exposure has been noted to repress SOCS expression through promoter methylation in human mast cells.^59^ Further, sustained exposure to IL-6 can upregulate IL6R in other cell types^60–62^ and is thus a plausible interpretation of the changes seen in iPSC-CMs. CXCR2 is known to rapidly undergo internalization in response to IL-8 binding;^63,64^ though little data exist regarding sustained IL-8 signaling in cardiomyocytes, increased expression at 24 hours of high-risk serum exposure may suggest receptor expression rebalancing. Importantly, theses interpretations remain inferential pending further mechanistic studies in iPSC-CMs.

Both IL-6 and IL-8 signaling can modulate JNK and the AP-1 transcription factor complex signaling,^65–68^ and high-risk serum led to a shift in AP-1 transcriptional factor subunit expression in iPSC-CMs with upregulation of JUN and downregulation of FOS, FOSB, and JUNB. Intriguingly, JNK and AP-1 signaling is active in cardiomyocytes^69^ and has been linked to sarcomere integrity and contractile function, as well as mitochondrial respiration. AP-1 subunit JUN deletion disrupted sarcomere structure in neonatal murine cardiomyocytes,^70^ and AP-1 inhibition depressed cardiomyocyte responses to beta-adrenergic stimulation or calcium supplementation.^71^ Inhibition of JNK phosphorylation impaired mitochondrial respiration during murine cardiomyocyte ischemia-reperfusion injury,^72^ although JNK signaling in cardiomyocytes and in mitochondria in particular is multifaceted.^73^ Ultimately, the role of JNK and AP-1 signaling in the human iPSC-CM response to risk-stratified septic serum requires direct mechanistic investigation.

In summary, we identified a serum proteomic shift across PERSEVERE-II mortality risk strata in children with septic shock and found that human iPSC-CM spheroids demonstrated differential functional and transcriptional responses to risk-stratified septic serum. High-risk serum exposure recapitulated features of clinical SAMD, including reversible depression of contractility, as well as mitochondrial dysfunction. We also identified distinct patterns of gene expression with high-risk serum and contextualized transcriptional changes with the proteomic identification of IL-6 and IL-8 as key differentially expressed proteins in children with high mortality probability. We believe that risk-stratified serum exposure to iPSC-CMs is an effective translational model of sepsis-associated myocardial dysfunction from which to investigate novel therapeutic approaches.

## ACKNOWLEDGMENTS

The authors are indebted to the mentorship of Dr. Hector R. Wong. The authors are grateful for technical expertise provided by Patrick Lahni and Adam Veteto. RNA sequencing was conducted with the assistance of the CCHMC Genomics Sequencing Facility. The authors are grateful to Amy Pitstick and the CCHMC Pluripotent Stem Cell Facility for technical support with iPSC culture and to Aditi Paranjpe and Ronika De with CCHMC Bioinformatics Collaborative Services regarding RNA sequencing analyses.

## SOURCES OF FUNDING

A.J.L. was supported by NIH K08GM148957. A.J.L., M.R.A., and B.Z. were supported by NIH R21GM150093. A.J.L., M.R.A., and B.Z. were supported by NIH R33GM151703. M.R.A. received funding through NIH R35GM155165 and a Procter K-to-R Scholar award through the Cincinnati Children’s Research Foundation.

## DISCLOSURES

None

---

**Supplemental Figure 1: Repeated measurements of iPSC-CM spheroids**. Percent shortening of individual iPSC-CM spheroids day+126 after differentiation (labeled A through K) over the course of three days. The median value for each spheroid is depicted as the horizontal bar. n=11 individual spheroids.

**Supplemental Figure 2: Proteomic analyses of serum from children with septic shock stratified by mortality risk**. Two-dimensional Principal Component Analysis plots of principal component (PC) 1 versus PC2 (**a**) and PC2 versus PC3 (**b**), to visualize differences in proteomes between low-, intermediate-, and high-risk strata. Proteomic shift is best visualized on the three-dimensional plot. Reactome pathway annotation of under-represented proteins (**c**), as well as Gene Ontology Biological Process pathway annotation of over-expressed (**d**) and under-expressed (**e**) proteins in high-risk children compared to low-risk children with septic shock with a Benjamini-Hochberg adjusted p-value < 0.05 after adjustment for plate, age, and site.

**Supplemental Figure 3: iPSC-CM spheroid contractility**. Video clip of iPSC-CM spheroid with spontaneous contractions.

## Notes

### Competing Interest Statement

The authors have declared no competing interest.

